## Supporting figures and legends for "Nanopore sequencing reveals that DNA replication compartmentalisation dictates genome stability and instability in *Trypanosoma brucei*"

**Supplementary figure legends**

**Table S1. Nanopore sequencing of the *T. brucei* Lister 427 genome.**

**Table S2. Comparing genome assembly of the Lister 427 *T. brucei* genome by Nanopore and Pacbio/HiC analysis, and of *T. brucei* TREU927.**

**Fig.S1. Nanopore contigs greater than 1 Mb in size.** 13 Nanopore contigs (tigs) are shown in which core genes are shown organised as directional gene clusters (dark grey and light depicting the transcribed strand), centromeres are shown in blue, and *VSG*s within the subtelomeres in red.

**Table S3. Summary of Nanopore assembly of the 11 *T. brucei* megabase chromosomes.**

**Fig.S2. Connecting the cores, subtelomeres and VSG expression sites of the *T. brucei* megabase chromosomes through Nanopore long-read sequencing.** Circos plots highlighting synteny between the Muller genome assembly and Nanopore contigs (tigs) of chromosome 10 (A), chromosome 4 (B), and three further chromosomes (C). In all panels grey ribbons represent overlaps, black ribbons represent multiple overlaps within a region, red denotes *VSG* genes, and blue denotes a centromere; in each case, the transcribed core, transcriptionally silent subtelomeres (numbered 3A, 3B, 5A, 5B), and bloodstream VSG expression sites (BESS) in the Muller genome are labelled, with the organisation of these compartments diagrammed (adapted from [9]).

**Fig.S3. R-loop mapping across the megabase chromosomes and DNA repeat regions of the *T. brucei* genome. A.** Mapping of R-loops (DNA-RNA hybrid immunoprecipitation and sequencing, DRIP; data shown as IP/input) in wild type (WT) bloodstream from cells and RNase H1 null (-/-) mutants is shown in five Nanopore contigs (tigs) that span the core and subtelomere compartments. **B.** Metaplots of DRIP-seq signal in WT and RNase H1-/- cells across all Nanopore assembled regions containing 177 bp, 50 bp, 70 bp and centromeric (centro) repeats.

**Fig.S4. Distinct base composition of the core and subtelomeres of the *T. brucei* megabase chromosomes.** GC skew, AT skew and GC content is shown for four Nanopore contigs (tigs) that span the core (grey) and subtelomeric (red) compartments of *T. brucei* megabase chromosomes. Base skew is shown relative to transcribed strands of directional gene clusters (blue, forward strand; red, reverse strand).

**Fig.S5. Compartmentalisation of DNA replication between the cores and subtelomeres of the *T. brucei* megabase chromosomes.** The distribution of DNA replication origins across the core and associated subtelomeres of the 11 megabase chromosomes, assessed by MFA-seq , is shown using the Muller genome (tritrypDB, release v46) as a reference. Directional gene clusters in each chromosome core (dark grey and light depicting the transcribed strand), centromeres (blue), and *VSG*s (red) within the subtelomeres are indicated. The graphs above each chromosome show the read depth ratio of Illumina sequence derived from DNA of the following cells: early S phase/G2-M phase cells (dark red, bloodstream cells, BSF; dark green, procyclic form, PCF), late S/G2M (light red, BSF; light green, PCF), and G1/G2M (grey for both life cycle stages); each point represents the median S/G2M or G1/G2 ratio (y-axis) in 1 kb bins across the contig. All graphs are scaled according to chromosome compartment size.

**Fig.S5. Compartmentalisation of DNA replication between the cores and subtelomeres of the *T. brucei* megabase chromosomes.** The distribution of DNA replication origins across the core and associated subtelomeres of the 11 megabase chromosomes, assessed by MFA-seq , is shown using the Muller genome (tritrypDB, release v46) as a reference. Directional gene clusters in each chromosome core (dark grey and light depicting the transcribed strand), centromeres (blue), and *VSG*s (red) within the subtelomeres are indicated. The graphs above each chromosome show the read depth ratio of Illumina sequence derived from DNA of the following cells: early S phase/G2-M phase cells (dark red, bloodstream cells, BSF; dark green, procyclic form, PCF), late S/G2M (light red, BSF; light green, PCF), and G1/G2M (grey for both life cycle stages); each point represents the median S/G2M or G1/G2 ratio (y-axis) in 1 kb bins across the contig. All graphs are scaled according to chromosome compartment size.

**Fig.S6. DNA replication initiation at all assembled *T. brucei* centromeres.** Metaplots and heatmaps of MFA-seq signal across centromere candidates and flanking sequence up to 200 kb upstream and downstream. BSF, bloodstream- form cells; PCF, procyclic form cells. Signal is the ratio of read depth in early S-phase, late S-phase or G1-phase cells relative to G2-M cells. In the heatmaps where there is not enough sequence data upstream or downstream, the region is coloured white.

**Table S4. Summary of Nanopore assembly of *T. brucei* centromeres.**

**Fig.S7. Compartmentalisation of stability between the cores and subtelomeres of the *T. brucei* megabase chromosomes.** Read depth mapping is shown across the core and subtelomeric compartments in two clones of wild type (WT), RAD51 null mutant and BRCA2 null mutant cells, as follows: as a population before growth (P0), as a population after 23 passages of growth in culture (P23), and in three clones (1, 2, 3) generated from the populations after 23 passages of growth; data is show as a heatmap of RPKM (reads per kb per million reads). The innermost circles show MFA-seq data as plotted in Fig.3.

**Table S5. Summary of Nanopore assembly of *T. brucei* VSG expression sites, including upstream 50 bp repeats and subtelomere sequence and inclusion of the telomere tract.**

**Fig.S8. 177 bp and 59 bp repeats are found in both the sub-megabase chromosomes and in the centromeres of the megabase chromosomes. A.** Organisation of 177 bp and 59 bp repeats in three Nanopore contigs (tigs) of sub-megabase chromosomes; the location of these features is shown relative to the genes within the contigs and a telomere. **B.** Summary of Nanopore contigs containing assembled centromeres in which 177 bp and 59 bp repeats can be detected; the number of the repeat motifs in each centromere is indicated. **C.** Two Nanopore contigs of parts of megabase chromosomes, showing the extent of 177 bp and 59 bp repeats within the putatively fully assembled centromeres; core genes are shown in grey and *VSG*s in red.

**Fig.S9. A comparison of transcript abundance derived from the sub-megabase chromosomes and from the core and subtelomere compartments of the megabase chromosomes.** A comparison of RNA-seq mapping to the non-repetitive components of eight sub-megabse chromosome Nanopore contigs (tigs, smaller chromosomes), excluding the VSG BESs, and to eight megabase chromosome Nanopre contigs that span the core and subtelomeric compartments; in all cases *VSG*s are in red and other genes in dark or light grey depending on transcription direction. RNA-seq mapping is shown as RPKM (reads per kb per million reads), and the blue line denotes 100 RPKM.

### Table S1

**SEQUENCING STATISTICS**

| | |
|---|---|
| Total number of reads sequenced | 319 075 |
| Total bases sequenced | 3 558 525 709 |
| Mean read length (bp) | 11 152.6 |
| Median read length (bp) | 4307 |
| Read length N50 | 28 458 |
| Longest reads (bp) | 345 688 |
| | 277 178 |
| | 252 902 |
| | 241 633 |
| | 240 202 |

### Table S2

| Genome statistics | NANOPORE 427 | 427 2018 | 427 |
|---|---|---|---|
| # contigs | 166 | 317 | 32 |
| # contigs (>= 10000 bp) | 104 | 303 | 32 |
| # contigs (>= 25000 bp) | 100 | 183 | 31 |
| # contigs (>= 50000 bp) | 97 | 72 | 24 |
| largest contig | 5080222 | 4633729 | 4977113 |
| total length | 55332974 | 50081021 | 26754408 |
| total length (>= 25000 bp) | 55128177 | 47671040 | 26743874 |
| total length (>= 50000 bp) | 54988747 | 43945746 | 26465291 |
| N50 | 2194184 | 1412180 | 2482252 |
| N75 | 538565 | 539199 | 1619978 |
| L50 | 9 | 11 | 4 |
| L75 | 22 | 25 | 7 |
| GC (%) | 43.85 | 43.71 | 46.71 |
| # N's | 0 | 49000 | 1050844 |
| # N's per 100 kbp | 0 | 97.84 | 3927.74 |
| # predicted genes (unique) | 20830 | 17122 | 9102 |
| # predicted genes (>= 0 bp) | 21404 + 2 part | 17733 + 9 part | 9325 + 3 part |
| # predicted genes (>= 300 bp) | 19253 + 2 part | 16098 + 8 part | 8676 + 3 part |
| # predicted genes (>= 1500 bp) | 5305 + 0 part | 5322 + 0 part | 3526 + 2 part |
| # predicted genes (>= 3000 bp) | 1342 + 0 part | 1534 + 0 part | 1123 + 0 part |
| # VSG genes | 3511 | 3524 | 387 |
| BUSCO score | C: 139 [S: 128, D: 11], F: 23, M: 93, n: 255 | C: 141 [S: 138, D: 3], F: 22, M: 92, n: 255 | C: 140 [S: 137, D: 3], F: 22, M: 93, n: 255 |

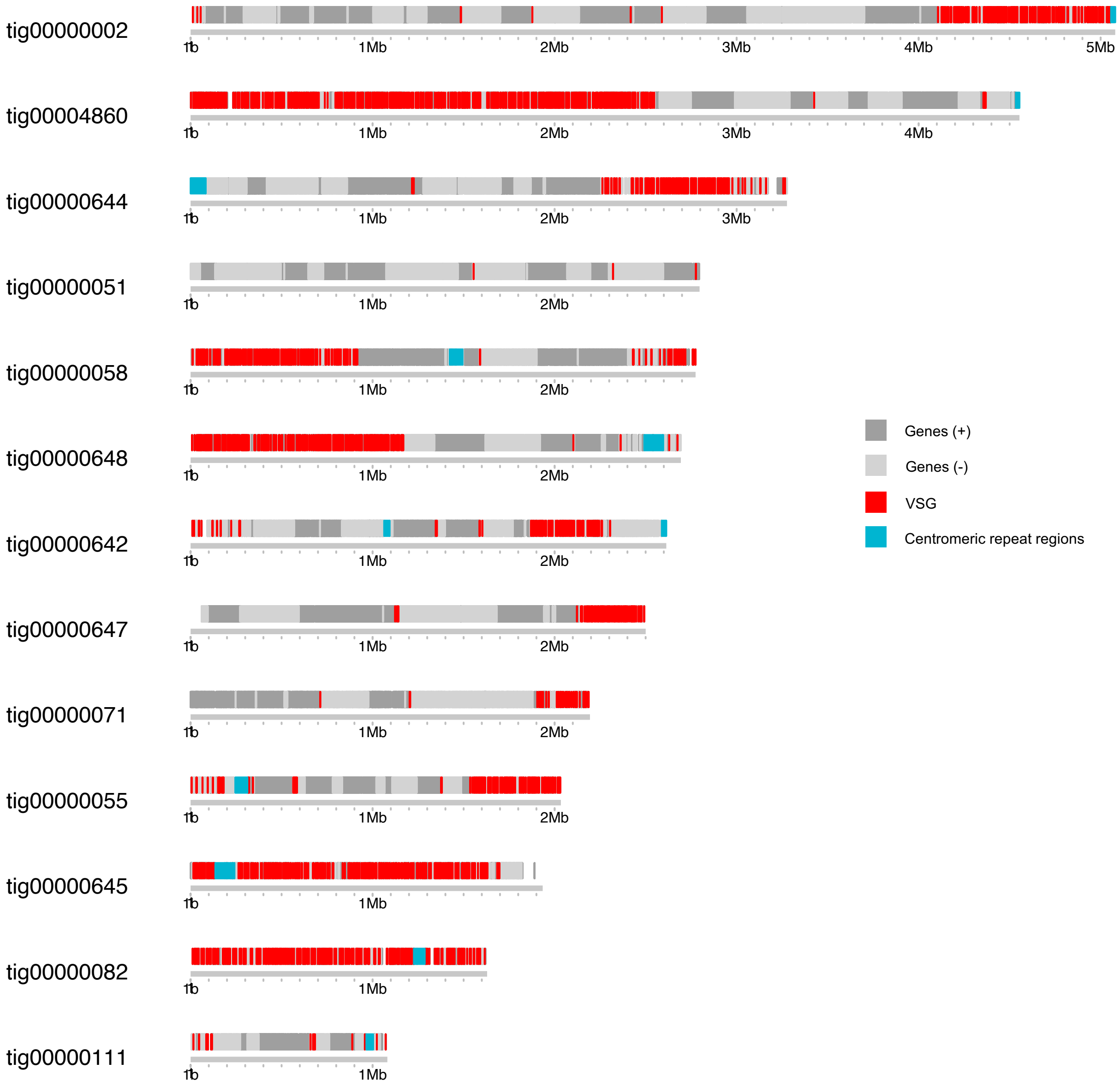

tig00000002
tig00004860
tig00000644
tig00000051
tig00000058
tig00000648
tig00000642
tig00000647
tig00000071
tig00000055
tig00000645
tig00000082
tig00000111
1b
1Mb
2Mb
3Mb
4Mb
5Mb
Genes (+)
Genes (-)
VSG
Centromeric repeat regions


Fig.S1

### Table S3

| Chr | Contigs within the reference | | | | | Contigs bridged in the new assembly |
|---|---|---|---|---|---|---|
| | **5A** | **5B** | **3A** | **3B** | **core** | |
| **1** | ✓ | ✓ | ✓ | ✓ | ✓ | 3A & core<br>3B & core<br>5B & core<br>5A & core |
| **2** | ✓ | | | | ✓ | |
| **3** | ✓ | ✓ | ✓ | | ✓ | 5A & core & 3A<br>5B & core |
| **4** | ✓ | ✓ | ✓ | ✓ | ✓ | 5A & 5B & core & 3A<br>3B & core |
| **5** | | | ✓ | ✓ | ✓ | 3A & core (& BES5)<br>3B & core |
| **6** | | | ✓ | ✓ | ✓ | 3B & core<br>3A & core |
| **7** | ✓ | | | | ✓ | 5A & core |
| **8** | ✓ | ✓ | ✓ | ✓ | ✓ | 5B & core<br>3A & core<br>5A & core |
| **9** | ✓ | ✓ | ✓ | ✓ | ✓ | 3B & core<br>3A & core<br>5A & core |
| **10** | ✓ | ✓ | ✓ | ✓ | ✓ | 3B & core<br>3A & core<br>5B & core |
| **11** | ✓ | ✓ | ✓ | ✓ | ✓ | 3B & core |

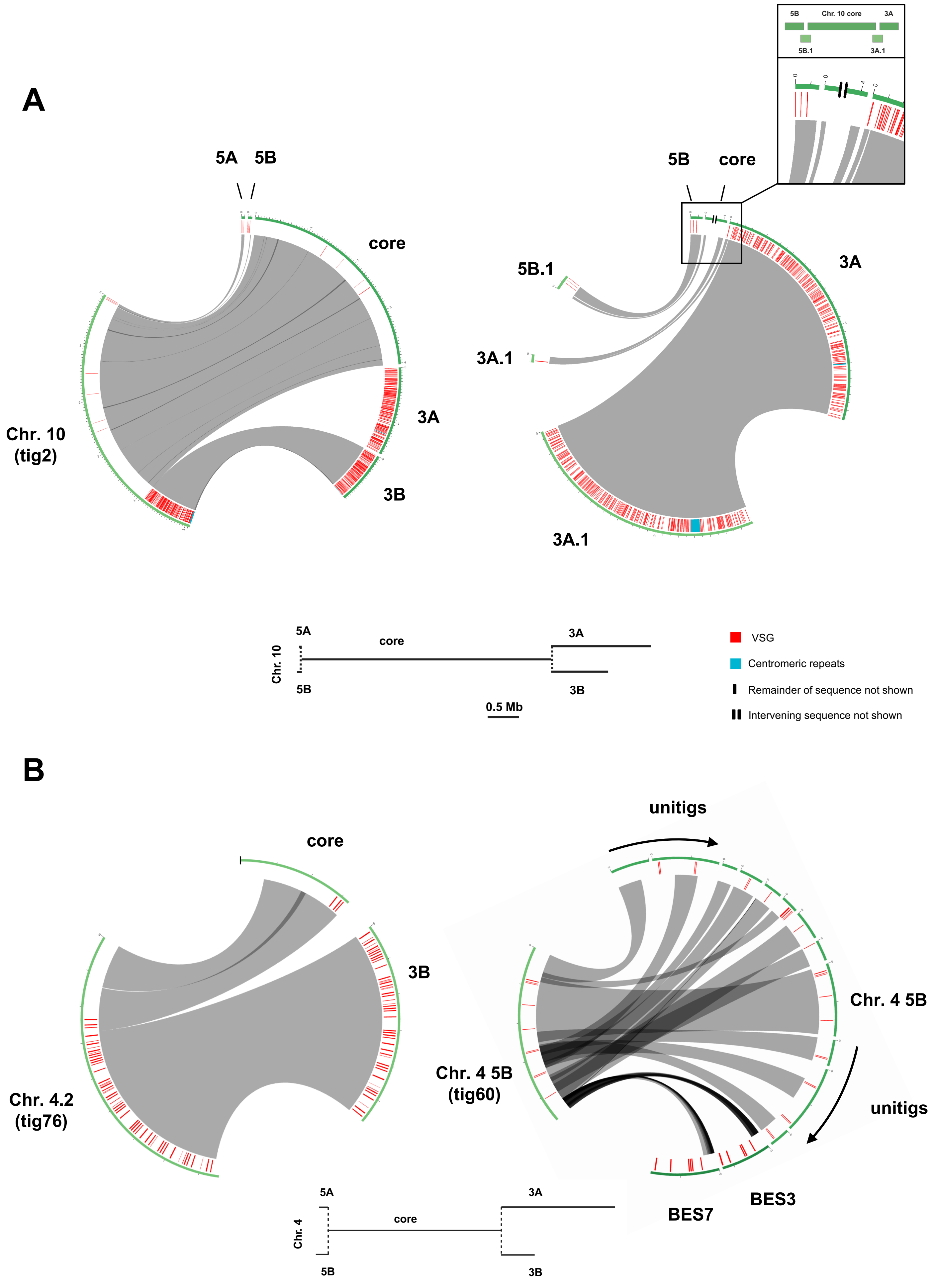
A
5A
5B
core
3A
3B
Chr. 10 (tig2)
5B
core
5B.1
3A
3A.1
3A.1
Chr. 10 core
5A
core
3A
Chr. 10
5B
3B
0.5 Mb
VSG
Centromeric repeats
Remainder of sequence not shown
Intervening sequence not shown
B
core
3B
Chr. 4.2 (tig76)
unitigs
Chr. 4 5B
Chr. 4 5B (tig60)
unitigs
BES3
BES7
5A
core
3A
Chr. 4
5B
3B
0.5 Mb

Fig.S2

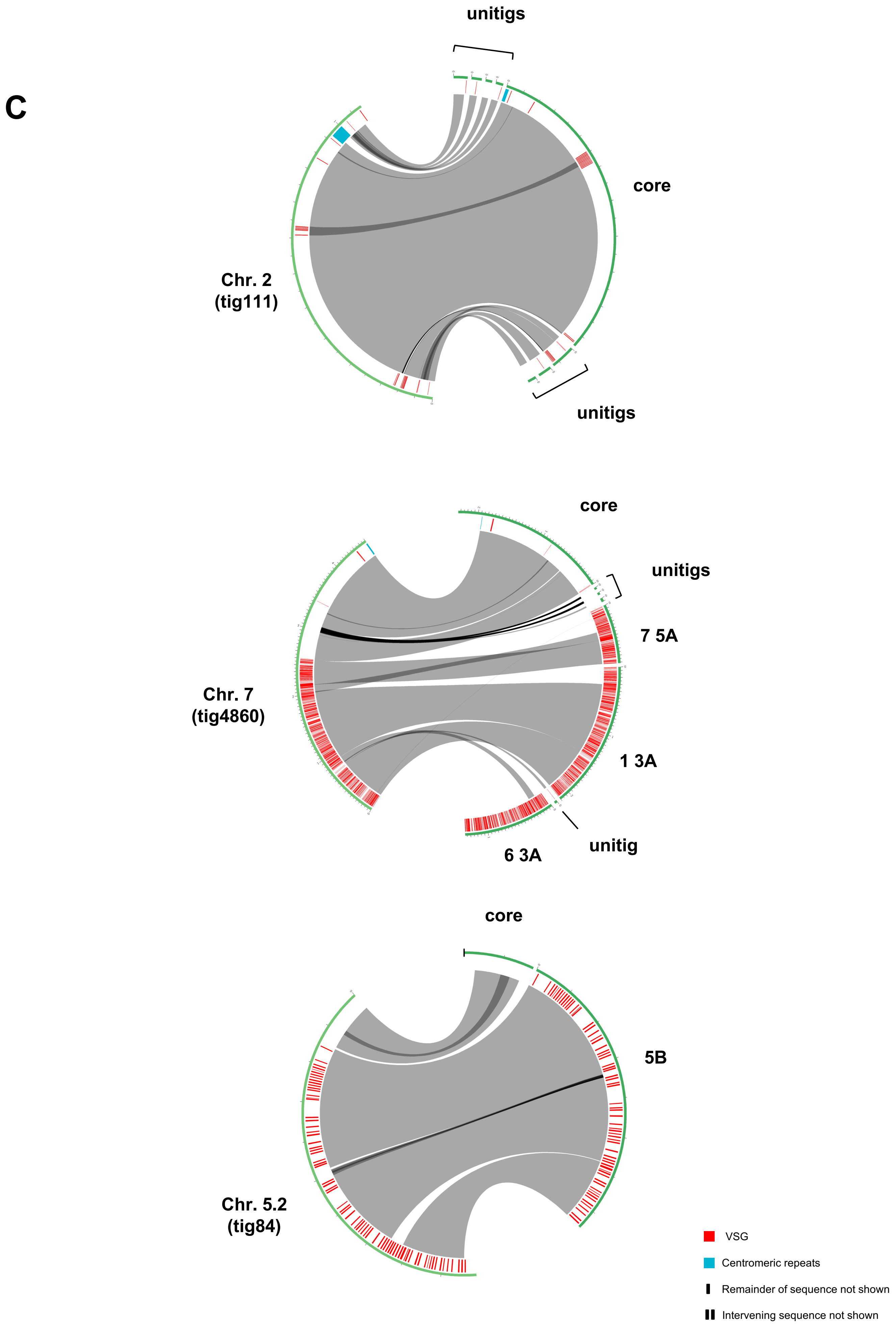


Fig.S2 continued

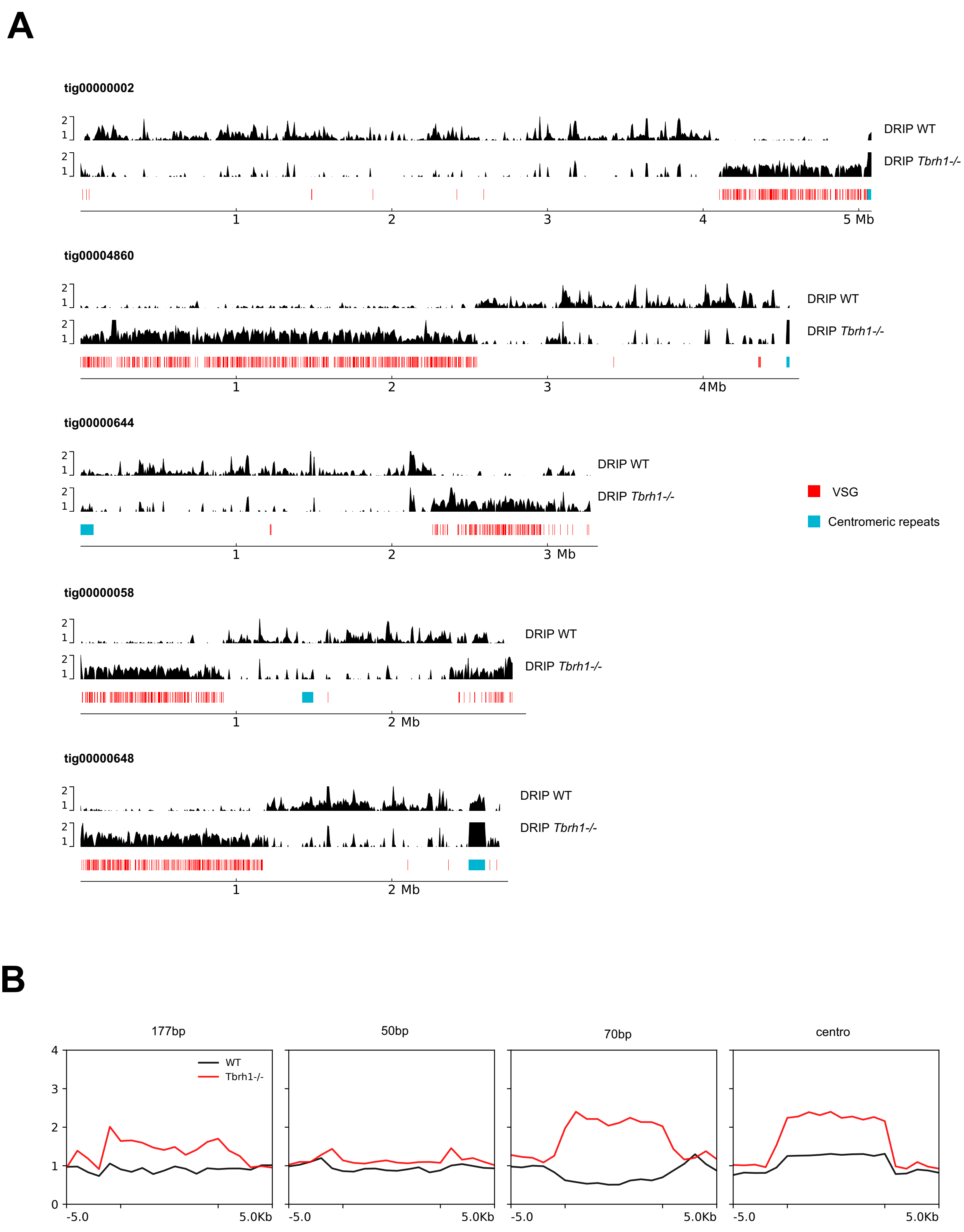
A
tig00000002
DRIP WT
DRIP *Tbrh1-/-*
1
2
3
4
5 Mb
tig00004860
DRIP WT
DRIP *Tbrh1-/-*
4Mb
tig00000644
DRIP WT
DRIP *Tbrh1-/-*
3 Mb
VSG
Centromeric repeats
tig00000058
DRIP WT
DRIP *Tbrh1-/-*
2 Mb
tig00000648
DRIP WT
DRIP *Tbrh1-/-*
2 Mb
B
177bp
50bp
70bp
centro
WT
Tbrh1-/-
-5.0
5.0Kb

Fig.S3

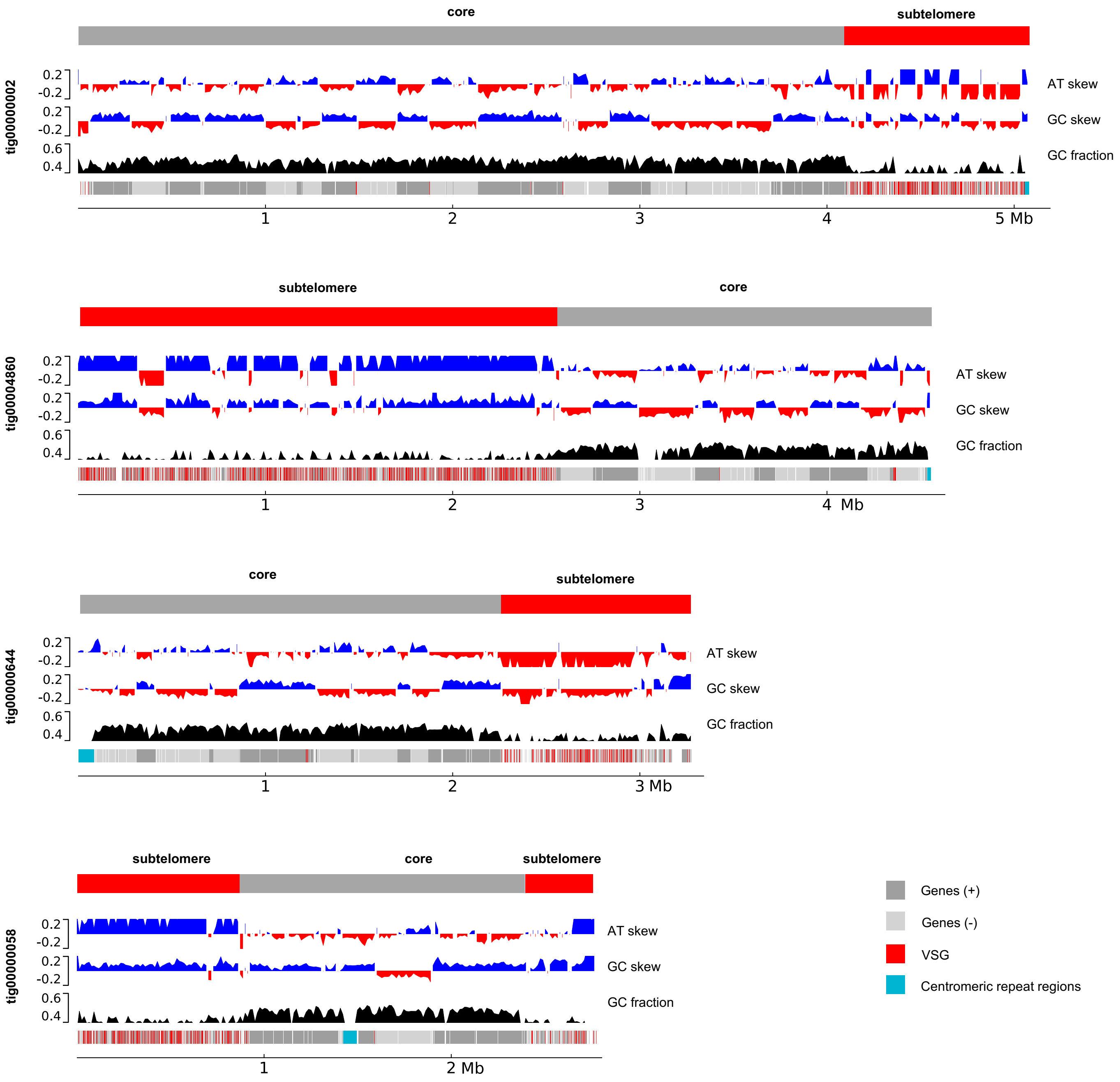
core
subtelomere
tig00000002
0.2
-0.2
0.2
-0.2
0.6
0.4
AT skew
GC skew
GC fraction
1
2
3
4
5 Mb
subtelomere
core
tig00004860
AT skew
GC skew
GC fraction
1
2
3
4 Mb
core
subtelomere
tig00000644
AT skew
GC skew
GC fraction
1
2
3 Mb
subtelomere
core
subtelomere
tig00000058
AT skew
GC skew
GC fraction
1
2 Mb
Genes (+)
Genes (-)
VSG
Centromeric repeat regions

Fig.S4

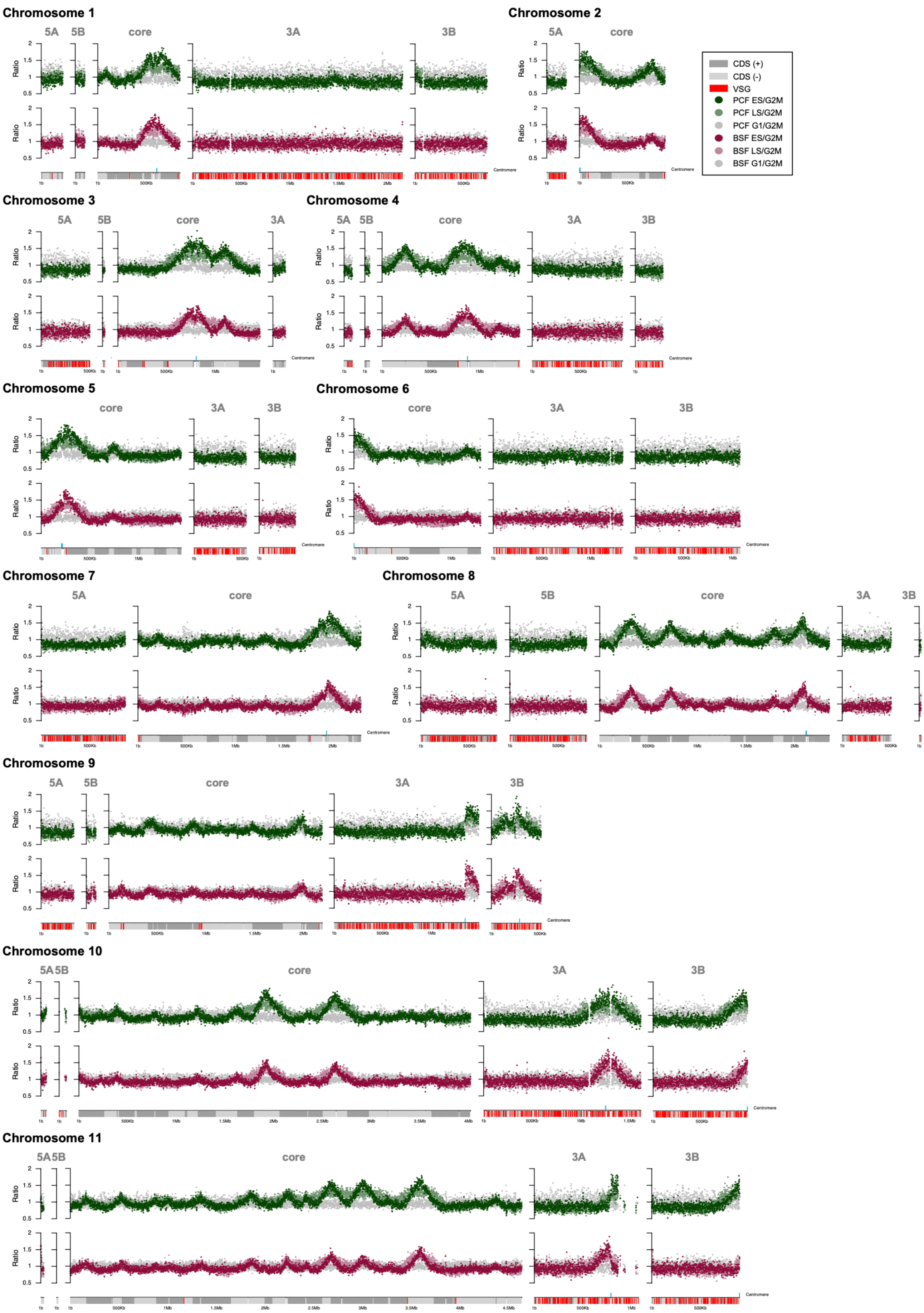

Chromosome 1
Chromosome 2
Chromosome 3
Chromosome 4
Chromosome 5
Chromosome 6
Chromosome 7
Chromosome 8
Chromosome 9
Chromosome 10
Chromosome 11
5A
5B
core
3A
3B
Ratio
Centromere
CDS (+)
CDS (-)
VSG
PCF ES/G2M
PCF LS/G2M
PCF G1/G2M
BSF ES/G2M
BSF LS/G2M
BSF G1/G2M


Fig.S5

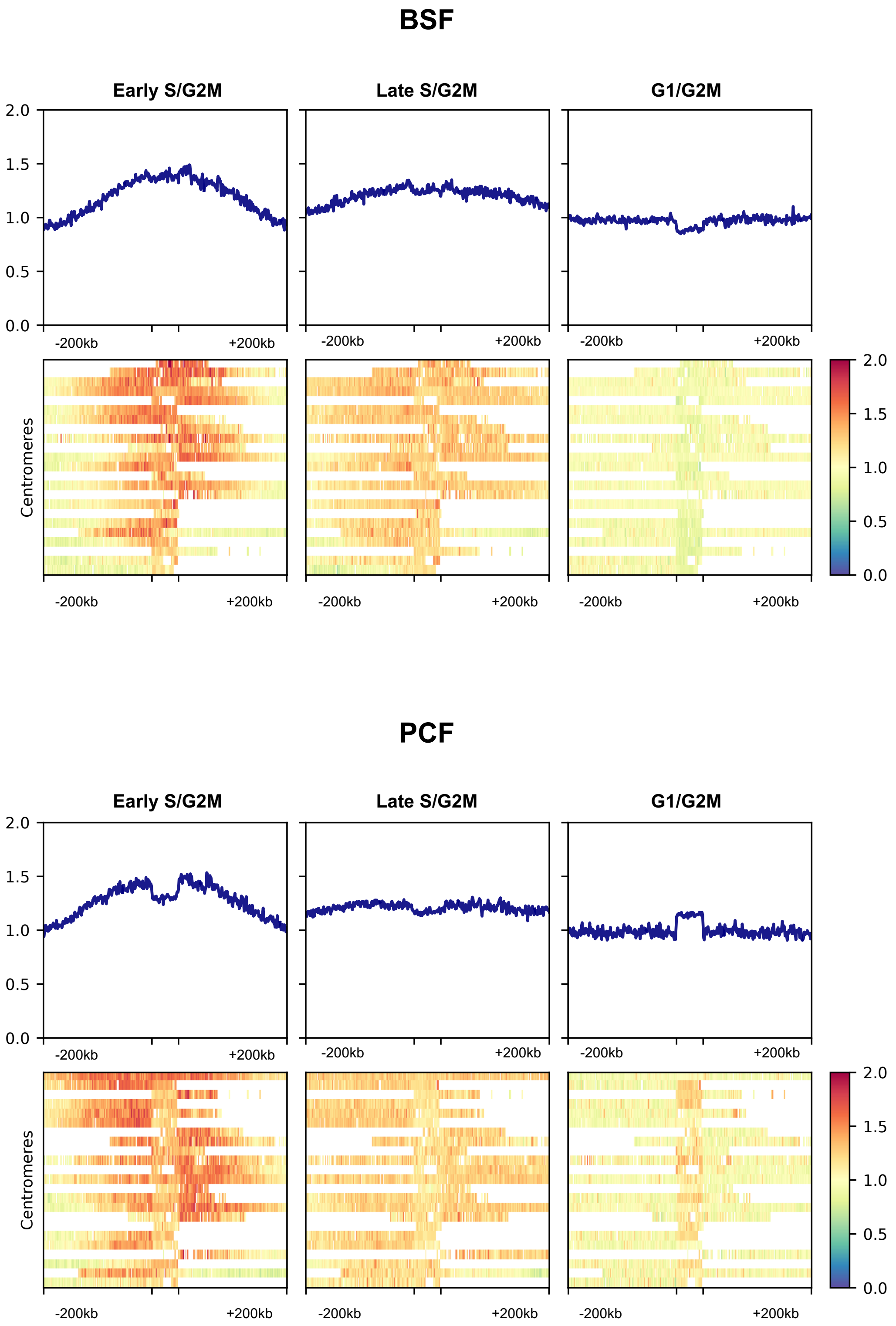
BSF
Early S/G2M
Late S/G2M
G1/G2M
2.0
1.5
1.0
0.5
0.0
-200kb
+200kb
Centromeres
PCF
Early S/G2M
Late S/G2M
G1/G2M
Centromeres

Fig.S6

### Table S4

| NO | CONTIG | CHR | SIZE (KB) | PUBLISHED SIZE (KB) | FULL-LENGTH | AT CONTENT (%) | PUBLISHED AT CONTENT (%) |
|---|---|---|---|---|---|---|---|
| **1** | tig00000654 | 1 | 20.4 | 20 & 65 | No | 70 | 66 |
| **2** | tig00000069 | 1 | 21.3 | | No | 69 | |
| **3** | tig00000111 | 2 | 37.2 | 30 & 55 | Yes | 73 | 66 |
| **4** | tig00000126 | 2 | 36.6 | | Yes | 72 | |
| **5** | tig00000642a | 3 | 30.2 | 75 & 80 | Yes | 50 | 49 |
| **6** | tig00000058 | 4 | 70.6 | 70 | Yes | 62 | 61 |
| **7** | tig00000055 | 5 | 66.5 | 50 & 75 | Yes | 62 | 62 |
| **8** | tig00000648 | 6 or 1 | 103.7 | 55 | Yes | 70 | 71 |
| **9** | tig00000127 | 7 or 1 | 17.0 | 100 & 120 | No | 73 | 75 |
| **10** | tig00004860 | 7 | 19.3 | | No | 73 | |
| **11** | tig00000032 | 8 | 39.5 | 100 | No | 62 | 59 |
| **12** | tig00000644 | 8 | 83.1 | | No | 62 | |
| **13** | tig00004861 | 8 | 44.4 | | No | 62 | |
| **14** | tig00000643 | 8 | 103.0 | | Yes | 61 | |
| **15** | tig00000642b | 8 | 24.5 | | No | 61 | |
| **16** | tig00000645 | 9 | 105.2 | Unknown | Yes | 71 | 60 |
| **17** | tig00000168 | 9 | 29.8 | | No | 68 | |
| **18** | tig00000002 | 10 | 22.9 | Unknown | No | 75 | 61 |
| **19** | tig00000082 | 10 | 59.5 | | Yes | 73 | |
| **20** | tig00000152 | 11 | 20.9 | Unknown | No | 71 | 61 |
| **21** | tig00000133 | 11 | 6.1 | | No | 70 | |
| **22** | tig00000115 | 2 or 7 | 14.6 | | No | 74 | |
| **23** | tig00000027 | unitig_2133 or 8 | 95.7 | | No | 62 | |

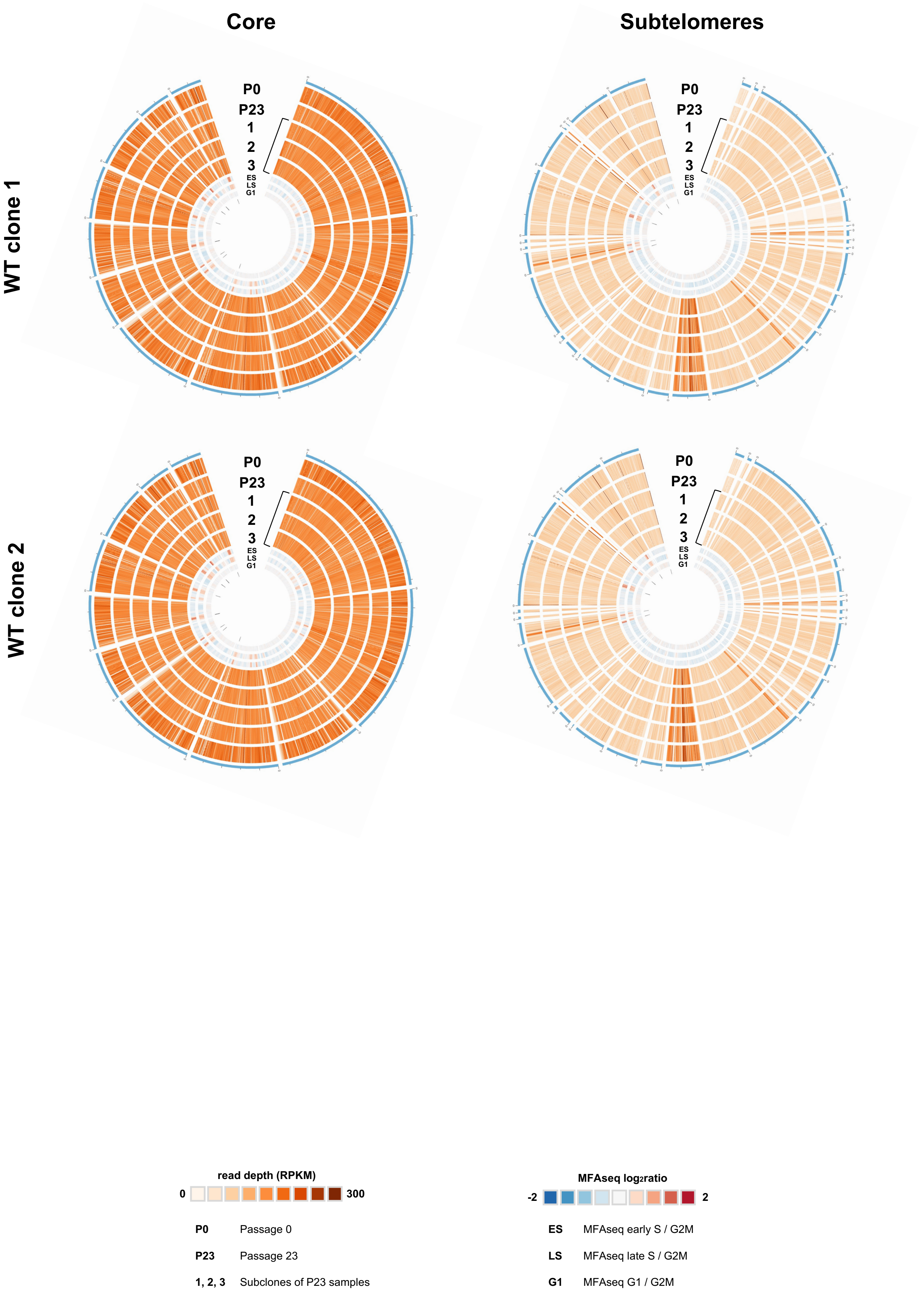
Core
Subtelomeres
WT clone 1
WT clone 2
P0
P23
1
2
3
ES
LS
G1
read depth (RPKM)
0
300
MFAseq log2ratio
-2
2
P0 Passage 0
P23 Passage 23
1, 2, 3 Subclones of P23 samples
ES MFAseq early S / G2M
LS MFAseq late S / G2M
G1 MFAseq G1 / G2M

Fig.S7

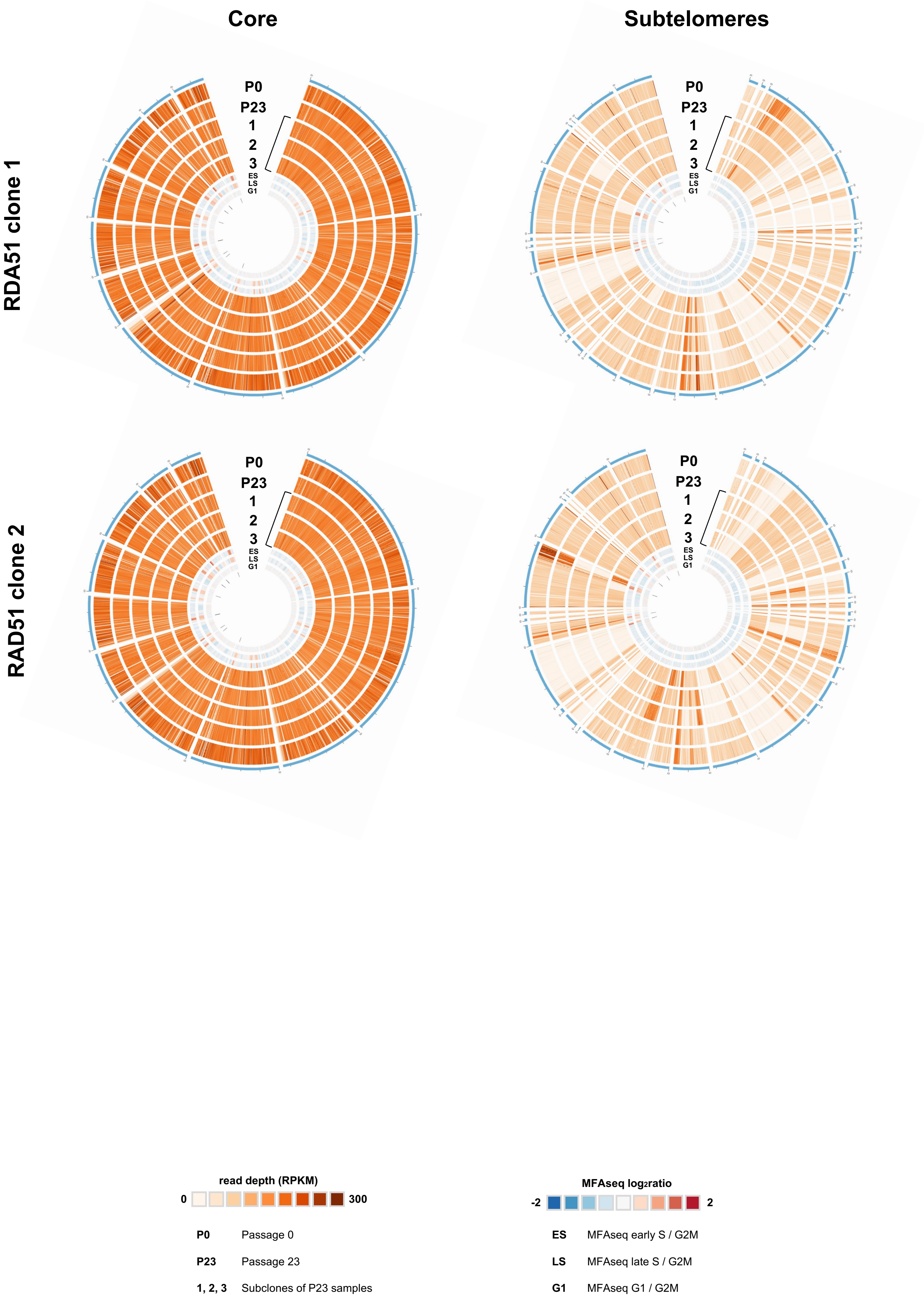


Fig.S7 continued

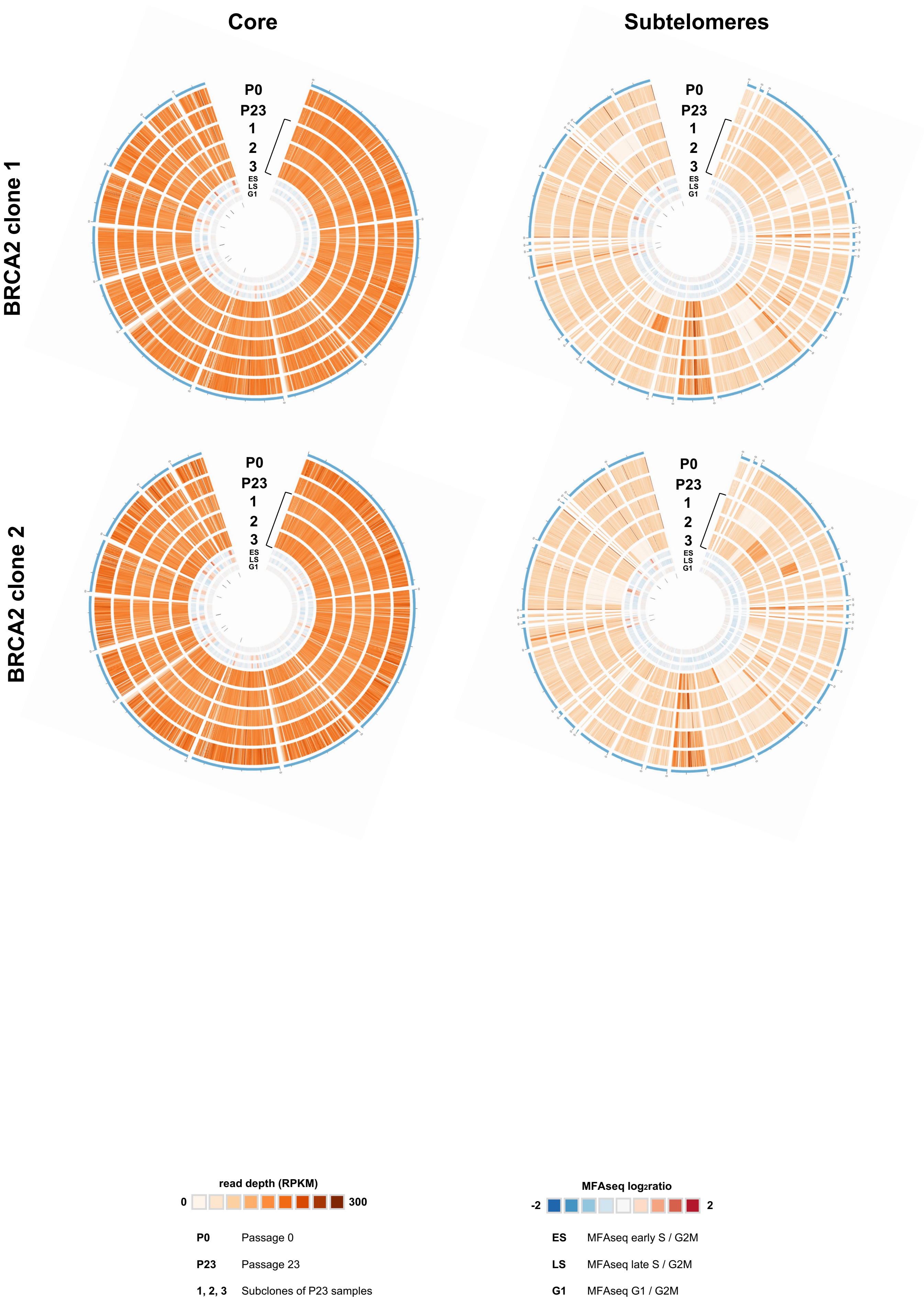


Fig.S7 continued

### Table S5

| NO | CONTIG | UP | 50BP | BES BODY | 70BP | CTR | BES VSG | TTAGGG | ID | CHR | CHR PUB | NOTES ON THE BES OR CONTIG |
|---|---|---|---|---|---|---|---|---|---|---|---|---|
| 1 | tig00000137 | ✓ | ✓ | ✓ | | ✓ | ✓ | ✓ | BES10 | 7 or int | int | no 70bp repeats |
| 2 | tig00000644 | ✓ | ✓ | ✓ | ✓ | ✓ | ✓ | ✓ | BES8? | 8 | 8 | very few genes in BES body |
| 3 | tig00000653 | | ✓ | ✓ | ✓ | ✓ | ✓ | ✓ | BES1 | | 6 | |
| 4 | tig00000114 | | ✓ | ✓ | ✓ | ✓ | ✓ | | BES12 | | 2 | |
| 5 | tig00000642 | ✓ | ✓ | ✓ | ✓ | ✓ | ✓ | | BES15 | 3 | 3 | two 70bp repeat regions |
| 6 | tig00004876 | ✓ | ✓ | ✓ | ✓ | ✓ | | | BES11 | int | int | 177bp repeats |
| 7 | tig00004877 | | ✓ | ✓ | ✓ | ✓ | | | BES11 | | int | |
| 8 | tig00004878 | | ✓ | ✓ | ✓ | ✓ | | | BES13 | | int | |
| 9 | tig00000055 | ✓ | ✓ | ✓ | ✓ | | | | BES5 | 5 | 5 | |
| 10 | tig00000156 | ✓ | ✓ | ✓ | ✓ | | | | BES14 | | 7 | |
| 11 | tig00000157 | ✓ | ✓ | ✓ | ✓ | | | | BES4 | | int | |
| 12 | tig00000658 | ✓ | ✓ | ✓ | ✓ | | | | BES7 | int | 6 | 177bp repeats |
| 13 | tig00004879 | ✓ | ✓ | ✓ | ✓ | | | | BES13 | int | int | 177bp repeats and telomere at the 5' end |
| 14 | tig00000116 | | ✓ | ✓ | ✓ | | | | BES3 | | 4 | |
| 15 | tig00000652 | ✓ | ✓ | ✓ | | | | | BES1 | | 6 | |

A

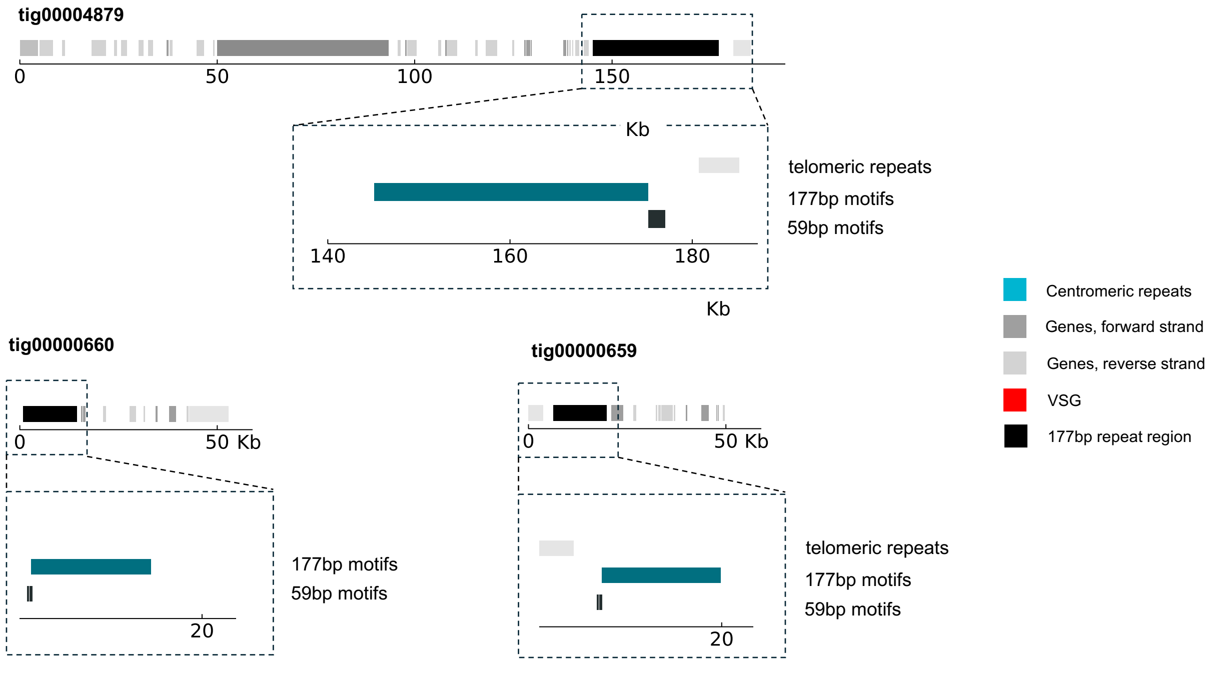


B

| TIG | CHR | 177BP | 59BP |
|---|---|---|---|
| **69** | 1 | 0 | 20 |
| **654** | 1 | 0 | 1 |
| **111** | 2 | 2 | 0 |
| **648** | 6 | 1307 | 244 |
| **645** | 9 (3A) | 51 | 0 |
| **168** | 9 (3B) | 58 | 200 |
| **82** | 10 (3A) | 284 | 5 |
| **2** | 10 (3B) | 1 | 0 |
| **152** | 11 (3A) | 59 | 404 |
| **4875** | 11 (3A) | 3 | 107 |
| **133** | 11 (3B) | 5 | 28 |

C

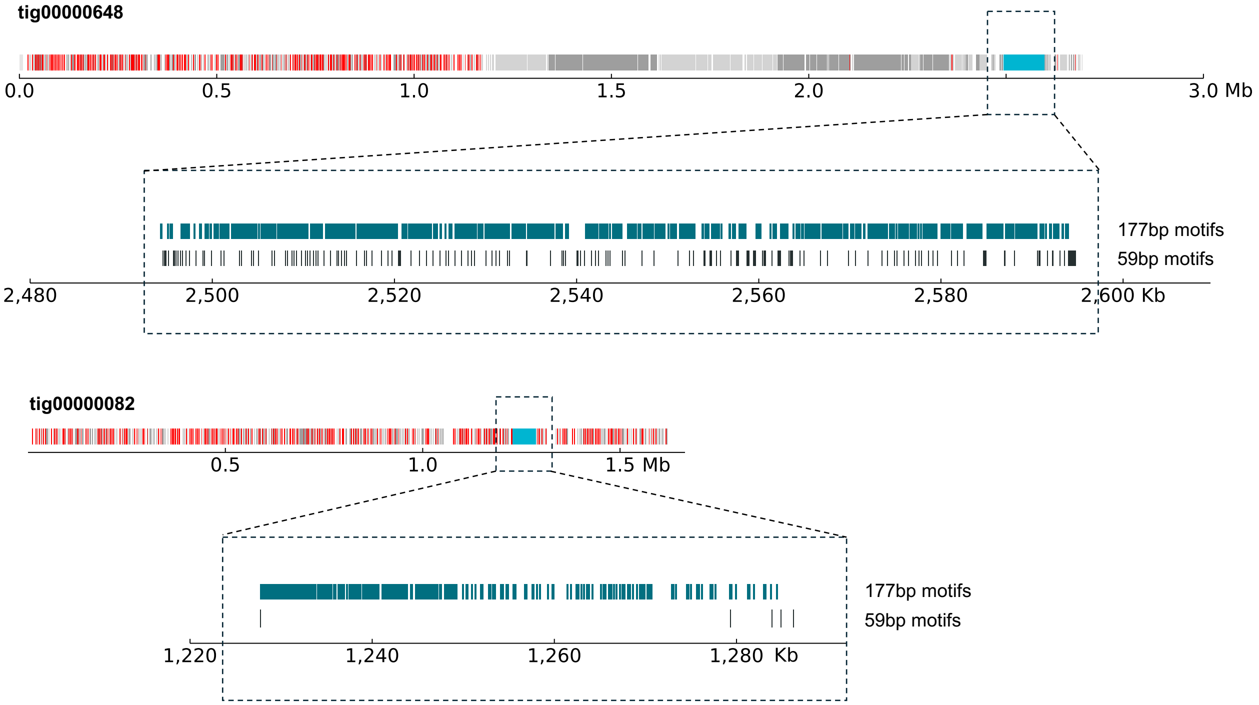


Fig.S8

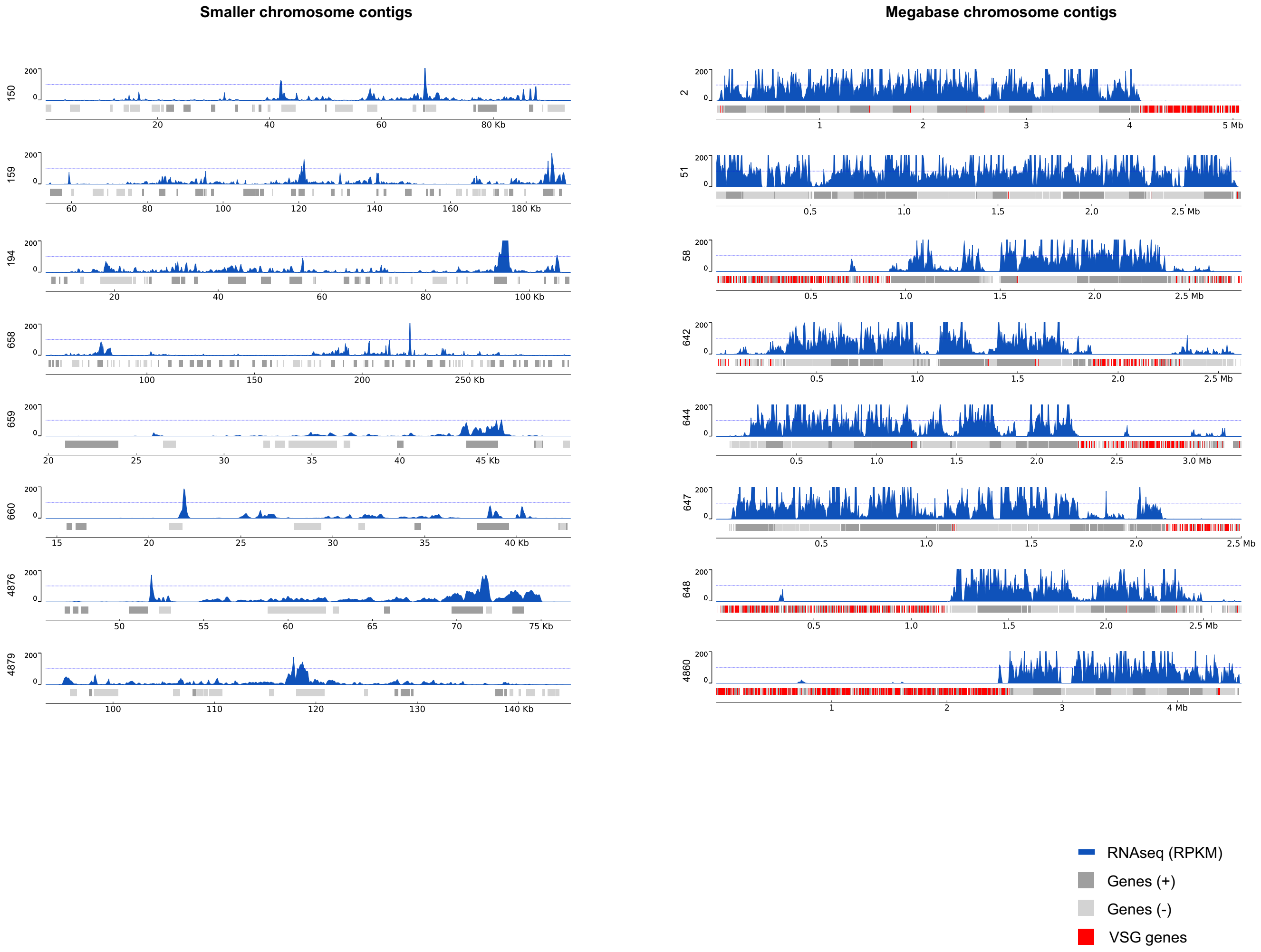
Smaller chromosome contigs
Megabase chromosome contigs
RNAseq (RPKM)
Genes (+)
Genes (-)
VSG genes

Fig.S9
